## Supporting information for "DDRP: real-time phenology and climatic suitability modeling of invasive insects"

**S1 Appendix. Estimating phenology model parameters for *Epiphyas postvittana*.**

The life stage durations used for the DDRP phenology model for *E. postvittana* (light brown apple moth) were estimated using data from a study by Geier and Briese (1980) of the relationship between rate of development and temperature (°C) of immature stages (egg, larvae + pupae) reared on Shorey medium [1] (presented in Fig. 3, p. 138). Using a lower threshold of 7.0°C, this study estimated the duration of the egg, larval, and pupal stage as 131, 421 (female larvae raised on young apple leaves), 510 (female larvae raised on old apple leaves), and 132 degree-days Celsius (DDC), respectively (Table 1).

Data on the oviposition schedule of adult females of two cohorts [1] (presented in Fig. 4, p. 139) were used to estimate the duration of the adult stage, which we defined as emergence of moths through 50% egg laying. One cohort of adult females were raised on Shorey medium and bulk-mated them in sets of eight males and eight females, while the other cohort was reared on broadbean plants and paired individually with a single mate. Moth emergence though 50% oviposition for the two cohorts took an average of 71 DDC (Table 2).

To keep the *E. postvittana* model for DDRP consistent with the one at USPEST.ORG, we rounded the lower developmental temperature threshold presented by Geier and Briese (1980) [1] from 44.6°F to 45°F. We used an empirical conversion to adjust the life stage durations according to this slightly higher threshold. This involved using weather data from 14 U.S. localities (seven for developing the factor, and seven more to test it) where the species is at high risk of establishment (Tables 3 and 4). We used the estimate for egg to 50% egg laying [1] as the generation time.

On average, the percent difference in degree-days when using a lower threshold of 44.6°F versus 45 °F was 97.1% (Table 3). We therefore multiplied life stage durations by 97.1%, which resulted in 229, 734, 231, and 128 degree-days Fahrenheit (DDF) [equal to 127, 408, 128, and 71 DDC; Table 1]. The validation test revealed an average error of 7.2 DDF when using a conversion factor to adjust the duration of egg to 50% egg-laying across the seven validation sites (Table 4).

**Table 1.** Duration of life stages of *E. postvittana* in degree-days Celsius and Fahrenheit based on a lower threshold (Tlow) of 7.0°C (44.6°F) compared to a Tlow of 7.2°C (45°F).

| | Tlow = 7°C (44.6°F) | | Tlow = 7.2°C (45°F) | |
|---|---|---|---|---|
| Stage | DDs (°C) | DDs (°F) | DDs (°C) | DDs (°F) |
| Egg | 131 | 236 | 127 | 229 |
| Larva | 421 | 756 | 408 | 734 |
| Pupa | 132 | 238 | 128 | 231 |
| Adult | 74 | 133 | 71 | 128 |

**Table 2.** Oviposition schedule of two cohorts of *E. postvittana* in degree-days Celsius based on a lower threshold of 7.0°C (44.6°F).

| | Cohort 1 | Cohort 2 | Average |
|---|---|---|---|
| Time to 5% OV | 20 | 35 | 28 |
| Time to 50% OV | 60 | 87 | 74 |
| Time to 75% OV | 92 | 127 | 110 |
| Time to 95% OV | 170 | 190 | 180 |

**Table 3.** Empirical conversion of the duration of egg to 50% egg-laying (in degree-days Fahrenheit) from a lower threshold (Tlow) of 44.6°F to a Tlow of 45°F.

| Replicate | DD (44.6F) | Date | DD (45F) | Percent |
|---|---|---|---|---|
| Medford OR | 1374 | 06/25/12 | 1330 | 96.8 |
| Yakima WA | 1373 | 06/29/12 | 1335 | 97.2 |
| Oak Ridge TN | 1385 | 05/03/12 | 1350 | 97.4 |
| Salisbury MD | 1364 | 05/21/12 | 1327 | 97.3 |
| San Lois Obispo CA | 1363 | 04/28/12 | 1323 | 97.1 |
| Salinas CA | 1371 | 05/06/12 | 1328 | 96.9 |
| Sacramento CA | 1385 | 05/11/12 | 1344 | 97.0 |
| Average | 1374 | | 1334 | 97.1 |
| St. Dev. | 8.87 | | 9.82 | 0.002 |

**Table 4.** Validation test of a conversion factor developed to empirically convert the duration of egg to 50% egg-laying (in degree-days Fahrenheit) from a Tlow of 44.6°F to a Tlow of 45°F. Sites used for validation were different from the ones used for developing the conversion factor.

| Replicate | DD (44.6F) | Date | DD (45F) | DD (45) estimated | Percent | Error / Absolute error in DDs[a] |
|---|---|---|---|---|---|---|
| Corvallis OR | 1365 | 07/03/12 | 1317 | 1326 | 96.5 | 9 / 9 |
| Fortuna CA | 1370 | 06/04/12 | 1320 | 1330 | 96.4 | 10 / 10 |
| Olema Valley CA | 1365 | 06/16/12 | 1318 | 1326 | 96.6 | 8 / 8 |
| Santa Rosa CA | 1365 | 06/09/12 | 1320 | 1326 | 96.7 | 6 / 6 |
| Norfolk VA | 1377 | 05/03/12 | 1339 | 1337 | 97.2 | -2 / 2 |
| Wilmington NC | 1382 | 04/18/12 | 1348 | 1342 | 97.5 | -6 / 6 |
| Hilton Head SC | 1372 | 04/01/12 | 1339 | 1332 | 97.6 | -7 / 7 |
| Average | 1371 | | 1329 | 1331 | 96.9 | 2.5 / 6.6 |
| St. Dev. | 6.67 | | 12.8 | 6.5 | 0.005 | 7.2 / 2.7 |

[a]The difference in degree-days (Tlow = 45°F) between actual calculated values and values estimated using the conversion factor of 97.1%, developed using the seven sites in Table 3.

**S2 Appendix. Data sources for validating a DDRP phenology model for *Epiphyas postvittana.***

We used three monitoring data sets to validate predictions of the dates of first spring egg laying and generation length for light brown apple moth, *E. postvittana*. For each data set, we estimated the date of the peak in first spring flight from bar plots or line plots of moth count data. The first monitoring data set was collected by the U.S. Department of Agriculture's Animal and Plant Health Inspection Service (USDA APHIS) in Alameda, Contra Costa, Monterey, and San Francisco counties in California from 2007 to 2009. Moth count data were reported only on a monthly basis in bar plots, and the precise locations of the study were not reported. Given the coarse spatial and temporal resolution of this data set, we assessed only whether the range of DDRP predictions of first spring egg laying for each county included the month when the observed peak spring flight occurred. We calculated the average, minimum and maximum date of model predictions across the DDRP grid cells for each county.

The second monitoring data set was collected by the University of California Cooperative Extension in and around wholesale nurseries, apples, and berries in five regions of Santa Cruz and Monterey counties in 2012, 2013, and 2014 [1]. Moth count data were collected on a bi-weekly basis and reported in bar plots. We estimated that each region could potentially overlap with four DDRP grid cells (i.e. 8 × 8 km) based on a map of the study regions provided in the report (Fig. 1). We therefore calculated the average, minimum and maximum date of model predictions across the four grid cells for each region. Data for regions in which peak flights were indiscernible were excluded from analyses.

The third monitoring data set was collected by USDA APHIS for a study of population dynamics of *E. postvittana* on four different host plants in Salinas, California in 2019 and 2020. Moth count data were collected on a bi-weekly basis and were available in both raw and line plot format. Peaks in fall and spring flight were virtually identical across the four host plants, so we used single estimates for both. We extracted DDRP model predictions for a single grid cell that overlapped with the location of the study (Fig. 1).

### References

**1.** Tjosvold S. Current Trap Data for Light Brown Apple Moth in Santa Cruz and Monterey Counties. Data: University of California Cooperative Extension, Santa Cruz County [Internet]. Available from: https://ucanr.edu/sites/uccesc/files/157533.pdf

**Fig. 1.** Map depicting the location of the study areas where the second and third monitoring data sets used to validate *E. postvittana* model predictions were collected. For the second population monitoring data set, we used a map that depicted the location of the five study regions (see pg. 1 of [1]) to estimate which DDRP grid cells each region overlapped with (blue squares). Model predictions for the third data set were extracted from a single grid cell that overlapped with the study location in Salinas (blue triangle). Red lines represent major highways and roads, and grid cells are colored according to the predicted date of first spring egg laying (warmer colors are later dates).

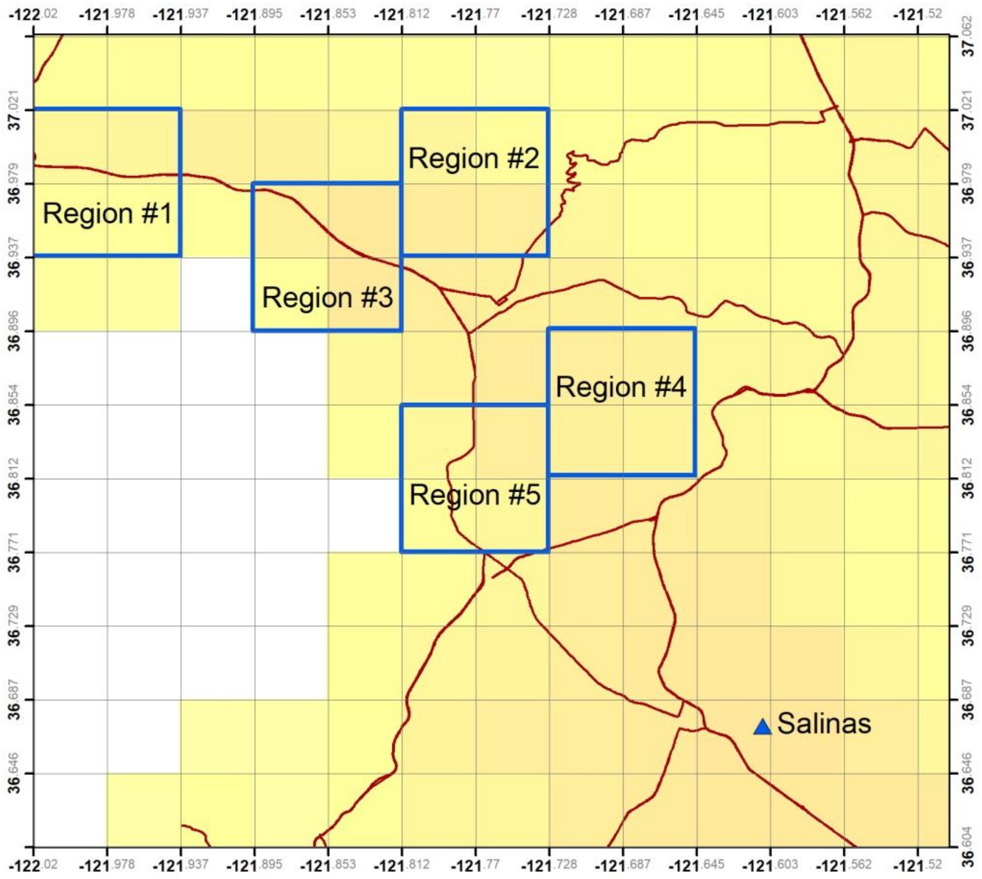

**S3 Appendix. Estimating a common lower temperature threshold and other phenology model parameters for *Neoleucinodes elegantalis.***

Moraes and Foerster (2015) [1] measured the development time of the immature stages of *Neoleucinodes elegantalis* at constant temperatures of 15, 20, 25, 27, and 30°C (50, 59, 68, 77 and 80.6°F; Table 1). They suggested a lower threshold of *ca*. 8.8°C for eggs, 7.7°C for larvae and pupae, and 17.5°C for pre-oviposition. Although large threshold disparities across stages can create problems for simple degree-day models that require a common threshold, the range of thresholds derived from this study was deemed to be a relatively minor issue. The authors did not test cooler temperatures [1], which lowers the accuracy of lower thresholds for each stage. Additionally, only three temperatures were used for measuring pre-oviposition, and the data point for 20°C is not well aligned with the other two points. This result suggests that the estimated lower threshold of 17.5°C may be too high.

We estimated a common lower threshold for *N. elegantalis* using a forced x-intercept method to re-analyze development times reported by Moraes and Foerster (2015) [1]. This involved adding a point to force the x-intercept to a common integer value in degrees Fahrenheit (48°F, 8.89°C) for the egg, egg-to-adult, and pre-oviposition (PreOV) stages (Table 1). Development rates and the development of each life stage in degree-day units were estimated using the regression equation for each stage (Table 2 and Figure 1). The larval and pupal (including prepupae) stage durations in degree-day units were estimated by subtracting the duration of the egg stage from the total egg-to-adult duration and multiplying the resulting value by the average proportion of days to development represented by each stage (Table 3). The summary of the phenology model is reported in Table 4.

In order to test the forced x-intercept approach against alternatives, we compared predictions for egg-to-adult development based on the forced x-intercept model to predictions based on 1) an unforced (i.e. unaltered) model and 2) an unforced model where the lower threshold was simply rounded to 48°F (Table 5). We found that forcing had little impact on predictions (Table 6). The values for the x-intercept and 1/slope for the egg-to-adult interval of *N. elegantalis* without forcing the regression were 8.64°C (47.6°F) and 1048.3, respectively. After constructing the model with forcing, these values were 8.89°C (48.0°F) and 1030.8, respectively. The use of the forced x-intercept method added an average of 1.2 days error compared to the unforced model; however, it had decreased error compared to the unforced model with a rounded threshold (average error = 2.5 days; Table 6).

**Table 1.** Development time in days for *N. elegantalis* reported by Moraes and Foerster (2015). An additional point (in bold font) was added to force the x-intercept to an integer value in degrees Fahrenheit.

| | | | Days Development | | | |
|---|---|---|---|---|---|---|
| Temp (°C) | Egg | Larvae | Prepupa | Pupa | Egg-to-Adult | PreOV |
| | **144** | | | | **580** | **164** |
| 15 | 15.5 | 40.4 | 9.1 | 31.4 | 96.4 | 9.8 |
| 20 | 7.3 | 25.6 | 4 | 13.2 | 50.1 | 6.5 |
| 25 | 5.3 | 15.9 | 3 | 9.4 | 33.6 | 3 |
| 27 | 4.9 | 16.7 | 2.6 | 9.2 | 33.4 | 3.8 |
| 30 | – | 13.2 | 2.1 | 8.8 | – | 4.3 |

**Table 2.** Development rates, lower threshold (Tlow), and stage durations in degree-days (DDs) for *N. elegantalis* using the modified x-intercept method. An additional point (in bold font) was added to force the x-intercept to an integer value in degrees Fahrenheit.

| | | | Development Rate (1/days) | | | |
|---|---|---|---|---|---|---|
| | Temp (°F) | Egg | Temp (°F) | Egg-to-Adult | Temp (°F) | PreOV |
| | **49** | 0.0069 | **50** | 0.0017 | **46.6** | 0.0061 |
| | 59 | 0.0645 | 59 | 0.0104 | 59 | 0.1020 |
| | 68 | 0.1370 | 68 | 0.0200 | 68 | 0.1538 |
| | 77 | 0.1887 | 77 | 0.0298 | 77 | 0.3333 |
| | 80.6 | 0.2041 | 80.6 | 0.0299 | 80.6 | 0.2632 |
| | 86 | – | 86 | – | | – |
| slope | (b) | 0.0064 | | 0.0010 | | 0.0102 |
| intercept | (a) | -0.3083 | | -0.0466 | | -0.4878 |
| $R^2$ | | 0.9948 | | 0.9902 | | 0.9216 |
| Tlow (°F) | (-a/b) | 48 | | 48 | | 48 |
| Tlow (°C) | (-a/b) | 8.9 | | 8.9 | | 8.9 |
| DDs (°F) | 1/slope | 156 | | 1031 | | 98 |
| DDs (°C) | 1/slope | 86 | | 573 | | 55 |

**Table 3.** Estimated proportion duration of larval and pupal (including prepupae) stages at three temperatures.

| | | Days Development | | | | Proportion | |
|---|---|---|---|---|---|---|---|
| Temp (°C) | Temp (°F) | Larvae | Pupae | Total | | Larvae | Pupae |
| 20 | 59 | 25.6 | 17.2 | 42.8 | | 0.60 | 0.40 |
| 25 | 68 | 15.9 | 12.4 | 28.3 | | 0.56 | 0.44 |
| 27 | 77 | 16.7 | 11.8 | 28.5 | | 0.59 | 0.41 |
| | | | | | Average | 0.58 | 0.42 |

**Table 4.** Summary of the degree-day model for *N. elegantalis* based on the forced x-intercept method.

| | Temp (°C) | Temp (°F) |
|---|---|---|
| Lower Threshold | 8.89 | 48 |
| Upper Threshold | 32.22 | 90 |
| Calculation Method | Single Sine | |
| Model Start | January 1$^{st}$ | |
| **Degree-Day Requirements** | **DDs (°C)** | **DDs (°F)** |
| Egg | 86 | 156 |
| Larva + pupa | 486 | 875 |
| Egg-to-Adult | 573 | 1031 |
| Pre-OV | 55 | 98 |
| DDs to Peak OV | 95 | 172 |
| DDs to 90% OV | 169 | 304 |
| Egg-to-1st-OV (min gen. time) | 627 | 1129 |
| Egg-to-Peak-OV (avg gen. time) | 668 | 1203 |
| **Events Summary** | **DDs (°C)** | **DDs (°F)** |
| First Spring Egg-Laying | 55 | 98 |
| Peak Spring Egg-Laying | 95 | 172 |
| First adults Gen. 1 | 627 | 1129 |
| Peak 1st Gen. Egg-Laying | 764 | 1375 |
| Peak 2nd Gen. Egg-Laying | 1432 | 2577 |
| Peak 3rd Gen. Egg-Laying | 2100 | 3780 |
| Peak 4th Gen. Egg-Laying | 2768 | 4983 |
| Peak 5th Gen. Egg-Laying | 3436 | 6186 |

**Table 5.** Development rate, lower threshold (Tlow), and duration in degree-days (DDs) for the egg-to-adult stage in *N. elegantalis* estimated using a forced and unforced model.

| | | Forced | Unforced |
|---|---|---|---|
| Temp (°F) | Days Develop. | 1/days | 1/days |
| 50 | 580.0 | 0.00172 | – |
| 59 | 96.4 | 0.01037 | 0.01037 |
| 68 | 50.1 | 0.01996 | 0.01996 |
| 77 | 33.6 | 0.02976 | 0.02976 |
| 80.6 | 33.4 | 0.02994 | 0.02994 |
| slope (b) | | 0.00097 | 0.00095 |
| intercept (a) | | -0.04657 | -0.04537 |
| $R^2$ | | 0.99022 | 0.97779 |
| Tlow (°F) | | 48.00 | 47.555 |
| Tlow (°C) | | 8.890 | 8.642 |
| DDs (°F) (1/slope) | | 1031 | 1048 |
| DDs (°C) (1/slope) | | 573 | 582 |

**Table 6.** Comparison of predicted dates for egg-to-adult development based on a forced model, unforced model, and unforced model with a rounded lower threshold (Tlow). Predictions were generated using three start dates for two generations (without inclusion of pre-oviposition times).

| | Forced | Unforced | Unforced with rounded threshold |
|---|---|---|---|
| Tlow (°F) | 48 | 47.56 | 48 |
| DDs (°F) | 1031 | 1048 | 1048 |
| Start date: 01/01/20 | | | |
| 1 generation | 07/03/20 | 07/02/20 | 07/04/20 |
| 2 generations | 08/22/20 | 08/20/20 | 08/23/20 |
| Start date: 04/01/19 | | | |
| 1 generation | 07/07/19 | 07/06/19 | 07/08/19 |
| 2 generations | 08/28/19 | 08/27/19 | 08/29/19 |
| Start date: 04/01/18 | | | |
| 1 generation | 07/08/18 | 07/07/18 | 07/09/18 |
| 2 generations | 08/23/18 | 08/22/18 | 08/26/18 |
| Avg. error (days) | 1.2 | 0 (reference) | 2.5 |

**Figure 1.** Linear regressions of development rates (1/days) for the egg, egg-to-adult, and pre-oviposition (PreOV) stages of *N. elegantalis* at constant temperatures of 50, 59, 68, 77 and 80.6°F (15, 20, 25, 27, and 30°C).

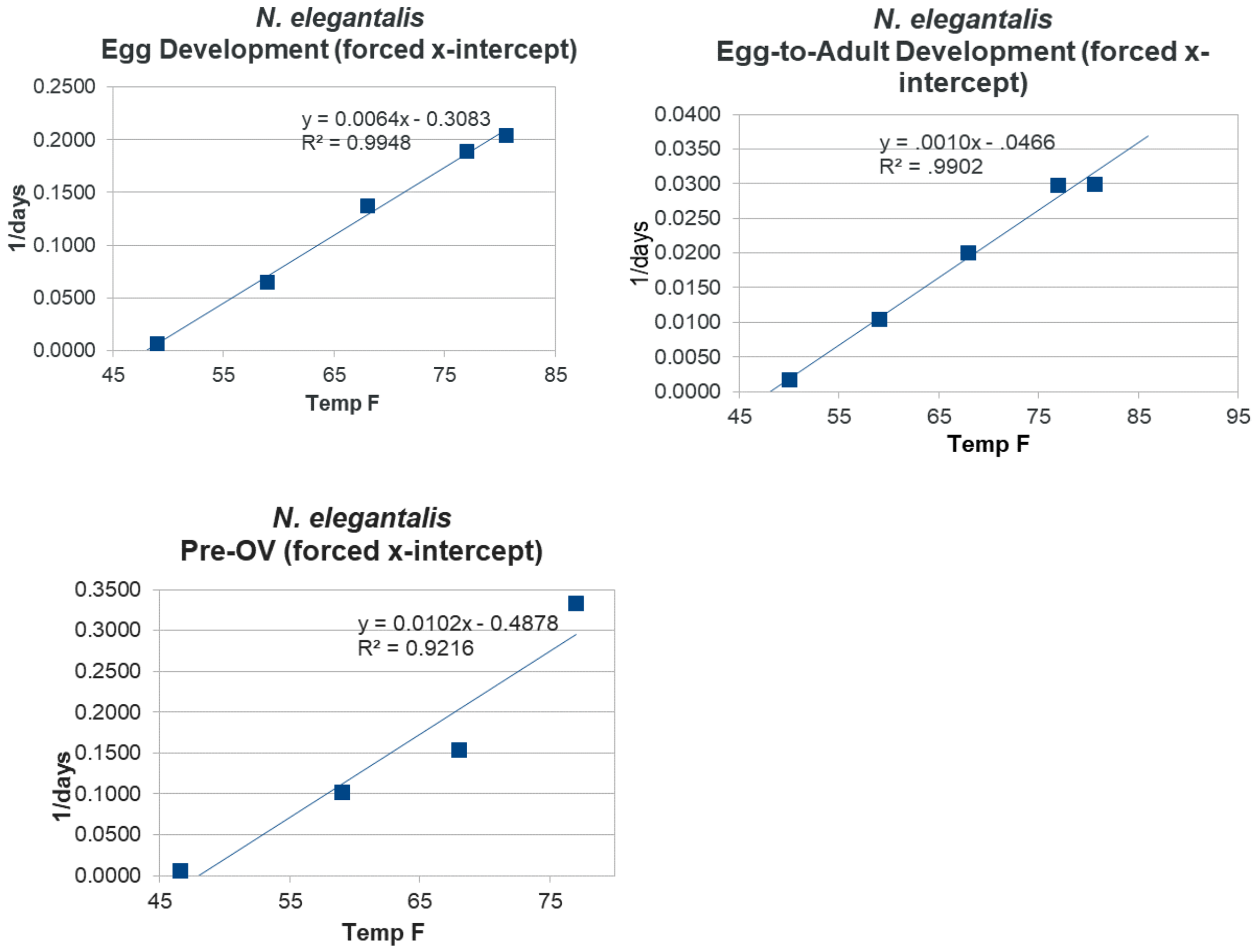

**S4 Appendix. Methods for fitting and validating a CLIMEX model for *Neoleucinodes elegantalis*.**

To fit and validate a CLIMEX model for *N. elegantalis*, we gathered 181 locality records from GBIF, the literature, graduate theses, and conference abstracts (S1 Table). We randomly subsampled 70% of the records (N = 127) for a model training set and reserved the remaining 30% of records (N = 54) for model validation. We added to the training data set an additional 101 localities that da Silva et al. (2018) used to validate their CLIMEX model for *N. elegantalis* [1], which resulted in a total of 228 localities for model fitting.

Our initial CLIMEX model applied the "best-fit" values presented by da Silva et al. (2018) [1], but this model excluded 22 model training localities from the potential distribution [Ecoclimatic Index (EI) = 0]. In particular, the model appeared to underpredict suitability for *N. elegantalis* in warmer areas of its known distribution. We therefore iteratively adjusted the limiting high temperature (DV3), heat stress temperature threshold (TTHS), and heat stress temperature rate (THHS) parameters and assessed how EI values changed in these areas. All CLIMEX simulations applied a top-up irrigation rate of 2.5 mm day$^{-1}$ for the winter and summer season. We found that the "high" values for these parameters that da Silva et al. (2018) included in their sensitivity analysis (DV3 = 31, TTHS = 31, and THHS = 0.00084) resulted in a good model fit. Specifically, all but six training localities in warmer areas had an EI > 0.

Additionally, we tested different cold stress threshold (TTCS) and cold stress temperature rate (THCS) values. To approximate the lowest temperatures that *N. elegantalis* may be exposed to, we extracted estimates of the historical (1950−2000) minimum temperature of the coldest week at a 10′ minute resolution (Bio6 in the CliMond v1.2 database [2]) for each training locality. The vast majority of localities (225/228 = 98.7%) did not experience historical monthly temperatures lower than 6°C, which suggests that cold stress accumulation at temperatures lower than 6°C may hinder establishment. We applied a TTCS and THCS of 6°C and −0. 0.0005, respectively.

**S1 Table. Locality records used for validating the CLIMEX and DDRP climatic suitability models for *Neoleucinodes elegantalis.*** The two-letter country code, locality name (if known), state or province, latitude, longitude, and reference number for each record is indicated. The "Approx." column indicates whether geographic coordinates were approximated from specific location information such as a city name (Approx. = 1), or if they were reported by the study or database (Apprx. = 0).

| Country | Locality | State/Province | Latitude | Longitude | Approx. | Reference |
|---|---|---|---|---|---|---|
| AR | Posadas | Misiones | -27.233 | -55.567 | 0 | 1 |
| AR | Orán | Salta | -22.871 | -64.363 | 0 | 2 |
| BR | | Algoas | -10.371 | -36.995 | 0 | 3 |
| BR | Manaus | Amazonas | -3.120 | -60.022 | 1 | 4 |
| BR | Empresa Brasileira de Pesquisa Agropecuaria | Brasília | -15.462 | -47.576 | 1 | 5 |
| BR | Jardim Botanico de Brasília | Brasília | -15.862 | -47.830 | 1 | 5 |
| BR | Reserva Ecologica do IBGE | Brasília | -15.949 | -47.878 | 1 | 5 |
| BR | Camocim de São Félix | Ceará | -8.347 | -35.774 | 1 | 6 |
| BR | Croatá | Ceará | -4.400 | -40.900 | 0 | 7 |
| BR | Guaraciaba do Norte | Ceará | -4.170 | -40.730 | 0 | 7 |
| BR | Jaburuna, Ubajara | Ceará | -3.854 | -40.921 | 0 | 8 |
| BR | Tiangua | Ceará | -3.734 | -40.996 | 1 | 9 |
| BR | Alegre | Espírito Santo | -20.752 | -41.489 | 0 | 10 |
| BR | Domingos Martins | Espírito Santo | -20.364 | -40.658 | 1 | 11 |
| BR | Vargem Alta | Espírito Santo | -20.673 | -41.010 | 1 | 11 |
| BR | Fazenda Agua Limpa | Goiás | -15.330 | -47.416 | 0 | 12 |
| BR | Fazenda Andreia, Abadia de Goiás | Goiás | -16.836 | -49.456 | 0 | 13 |
| BR | Fazenda Experimental da empresa Unilever, Goiaia | Goiás | -16.718 | -49.414 | 0 | 13 |
| BR | Fazenda Santa Rosa das Flores, Palminopolis | Goiás | -16.766 | -50.106 | 0 | 13 |
| BR | Goianápolis | Goiás | -16.505 | -49.021 | 1 | 14 |
| BR | Universidade Federal de Goiás, Goiania | Goiás | -16.600 | -49.278 | 1 | 15 |
| BR | Dourados | Mato Grosso do Sul | -22.23 | -54.802 | 1 | 16 |
| BR | Coimbra | Minas Gerais | -20.857 | -42.469 | 0 | 17 |
| BR | Rio Paranaiba | Minas Gerais | -19.223 | -46.205 | 0 | 18 |
| BR | Vicosa | Minas Gerais | -20.813 | -42.938 | 0 | 19 |
| BR | Rodovia Transamazonica (Altamira/Itaituba) | Pará | -3.463 | -52.895 | 1 | 20 |
| BR | São José dos Pinhais | Paraná | -25.535 | -49.206 | 0 | 21 |
| BR | Encruzilhada de São João | Pernambuco | -8.244 | -35.767 | 1 | 6 |

| Country | Locality | State/Province | Latitude | Longitude | Approx. | Reference |
|---|---|---|---|---|---|---|
| BR | Garanhuns | Pernambuco | -8.883 | -36.496 | 1 | 6 |
| BR | Petrolina | Pernambuco | -9.383 | -40.503 | 1 | 6 |
| BR | Cha-Grande | Pernambuco | -8.233 | -35.462 | 0 | 22 |
| BR | Camocim de São Felix | Pernambuco | -8.359 | -35.762 | 0 | 23 |
| BR | Experimental Station of Vitória de Santo Antão | Pernambuco | -8.114 | -35.291 | 0 | 24 |
| BR | Serra Talhada | Pernambuco | -7.938 | -38.313 | 0 | 25 |
| BR | Centro de Pesquisa Mokiti Okada, Ipeúna | Rio de Janeiro | -22.402 | -47.681 | 1 | 26 |
| BR | Federal Rural University of Rio de Janeiro, Seropedica | Rio de Janeiro | -22.750 | -43.683 | 0 | 27 |
| BR | Itaperuna | Rio de Janeiro | -21.199 | -41.892 | 1 | 28 |
| BR | | Rio de Janeiro | -22.451 | -42.771 | 0 | 29 |
| BR | | Rio Grande do Sul | -28.054 | -51.196 | 0 | 29 |
| BR | University of Santa Maria campus Frederico Westphalen | Rio Grande do Sul | -27.397 | -53.429 | 0 | 30 |
| BR | | Santa Catarina | -27.685 | -48.497 | 0 | 29 |
| BR | Experimental Station of Epagri, Lages | Santa Catarina | -27.800 | -50.317 | 0 | 31 |
| BR | São José do R. Pardo | São Paulo | -21.602 | -46.904 | 1 | 6 |
| BR | Divinolândia | São Paulo | -21.661 | -46.735 | 1 | 32 |
| BR | Taiúva | São Paulo | -21.124 | -48.454 | 1 | 32 |
| BR | Piracicaba | São Paulo | -22.823 | -47.757 | 0 | 33 |
| BR | Pongai | São Paulo | -21.733 | -49.367 | 1 | 34 |
| BR | Urupes | São Paulo | -21.202 | -49.294 | 1 | 35 |
| BR | Itabaiana | Sergipe | -10.689 | -37.432 | 1 | 36 |
| CO | El Peñol, La Piedra, El Porvenir | Antioquia | 6.222 | -75.179 | 0 | 2 |
| CO | El Peñol, Santa Inés, La Gabriela | Antioquia | 6.258 | -75.273 | 0 | 2 |
| CO | Jardín, La Linda, La Clara | Antioquia | 5.614 | -75.830 | 0 | 2 |
| CO | El Penol, Santa Ines | Antioquia | 6.257 | -75.271 | 0 | 37 |
| CO | Jardin, El Tapado/El Llano | Antioquia | 5.601 | -75.838 | 0 | 37 |
| CO | Santo Domingo | Antioquia | 6.477 | -75.137 | 0 | 37 |
| CO | Piedra Gorda | Antioquia | 6.240 | -75.480 | 1 | 38 |
| CO | San Rafael | Antioquia | 6.305 | -75.025 | 0 | 39 |
| CO | Buenavista, Patiño | Boyacá | 5.496 | -73.952 | 0 | 2 |
| CO | Anserma, Palo Blanco Alto | Caldas | 5.255 | -75.769 | 0 | 2 |
| CO | Anserma, San Pedro | Caldas | 5.256 | -75.797 | 0 | 2 |
| CO | Manizales, El Rosario | Caldas | 5.028 | -75.583 | 0 | 2 |
| CO | Manizales, San Peregrino | Caldas | 5.057 | -75.579 | 0 | 2 |
| CO | Palestina, La Parroquia | Caldas | 5.012 | -75.631 | 0 | 2 |

| Country | Locality | State/Province | Latitude | Longitude | Approx. | Reference |
|---|---|---|---|---|---|---|
| CO | Villa María, Tejares, Bachué | Caldas | 5.032 | -75.516 | 0 | 2 |
| CO | Reserva Ecologico Rio Blanco | Caldas | 5.126 | -75.447 | 0 | 29 |
| CO | Chinchina, San Andres/Los Alpes | Caldas | 4.934 | -75.614 | 0 | 37 |
| CO | Manizales, Bajo Tablazo Hoyo Frio/Los Eucaliptos | Caldas | 5.021 | -75.533 | 0 | 37 |
| CO | Inza, Belén | Cauca | 2.473 | -76.020 | 0 | 2 |
| CO | Inza, Pedregal | Cauca | 2.523 | -76.000 | 0 | 2 |
| CO | Nueva Estrella | Cordoba | 9.288 | -76.073 | 1 | 38 |
| CO | Fusagasugá, Espinalito, La Poderosa | Cundinamarca | 4.315 | -74.417 | 0 | 2 |
| CO | Fusagasugá, La Isla, El Recuerdo | Cundinamarca | 4.349 | -74.401 | 0 | 2 |
| CO | San Bernardo, Pirineos bajos, Buenavista | Cundinamarca | 4.151 | -74.426 | 0 | 2 |
| CO | Silvania, San Luis bajo | Cundinamarca | 4.384 | -74.630 | 0 | 2 |
| CO | Silvania, San Luís bajo, Villa | Cundinamarca | 4.704 | -74.618 | 0 | 2 |
| CO | Silvania, Santa Rita, Las Brisas | Cundinamarca | 4.409 | -74.361 | 0 | 2 |
| CO | Silvania, Victoria Alta | Cundinamarca | 4.424 | -74.357 | 0 | 2 |
| CO | Filandia, La Julia, Las Delicias | Departamento del Quindío | 4.698 | -75.682 | 0 | 2 |
| CO | Pijao, La Playa, El Billar | Departamento del Quindío | 4.335 | -75.712 | 0 | 2 |
| CO | Garzón, Fátima | Huila | 2.181 | -75.563 | 0 | 2 |
| CO | Gigante, Bajo Corozal | Huila | 2.339 | -75.516 | 0 | 2 |
| CO | Gigante, Tres Esquinas, La Ondina | Huila | 2.310 | -75.509 | 0 | 2 |
| CO | Santa Marta, Minca, El Campano | Magdalena | 11.124 | -74.098 | 0 | 2 |
| CO | Santa Marta, San Isidro, La Sirena | Magdalena | 11.186 | -74.009 | 0 | 2 |
| CO | San Pedro de Cartago | Nariño | 1.545 | -77.116 | 0 | 2 |
| CO | Arboleda | Nariño | 5.583 | -75.150 | 0 | 40 |
| CO | La Florida | Nariño | 1.302 | -77.411 | 0 | 40 |
| CO | La Union | Nariño | 1.605 | -77.134 | 0 | 40 |
| CO | San Pedro de Cartago | Nariño | 1.550 | -77.119 | 0 | 40 |
| CO | Tangua | Nariño | 1.095 | -77.394 | 0 | 40 |
| CO | Yacuanquer | Nariño | 1.115 | -77.401 | 0 | 40 |
| CO | Abrego, Llano Alto | Norte de Santander | 8.077 | -73.211 | 0 | 2 |
| CO | Abrego, Los Piñitos | Norte de Santander | 8.094 | -73.238 | 0 | 2 |
| CO | Ocaña, Llano Verde, Piedra gorda | Norte de Santander | 8.405 | -73.340 | 0 | 2 |
| CO | Ocaña, Llano Verde, Piedra gorda | Norte de Santander | 8.237 | -73.357 | 0 | 2 |
| CO | Pamplona, La Unión, La Morena | Norte de Santander | 7.264 | -72.577 | 0 | 2 |
| CO | Silos, Cherqueta, La Vega | Norte de Santander | 7.170 | -72.742 | 0 | 2 |
| CO | Dosquebradas, La Fría, Las Veraneras | Risaralda | 4.854 | -75.709 | 0 | 2 |

| Country | Locality | State/Province | Latitude | Longitude | Approx. | Reference |
|---|---|---|---|---|---|---|
| CO | Dosquebradas, La Fría, Las Veraneras | Risaralda | 4.855 | -75.710 | 0 | 2 |
| CO | Santa Rosa de Cabal, San Andrés, Los Alpes | Risaralda | 4.935 | -75.613 | 0 | 2 |
| CO | Sta Rosa de Cabal/El Embol | Risaralda | 4.920 | -75.633 | 0 | 2 |
| CO | Jesús María, Alto Grande, La Vega | Santander | 5.879 | -73.787 | 0 | 2 |
| CO | Mesa de los Santos, Acuarela | Santander | 6.759 | -73.106 | 0 | 2 |
| CO | Mesa de los Santos, Tabacal, El Alto | Santander | 6.792 | -73.056 | 0 | 2 |
| CO | Pie de Cuesta, Sevilla | Santander | 7.026 | -72.994 | 0 | 2 |
| CO | Cajamarca, Arenillal, La Esperanza | Tolima | 4.391 | -75.483 | 0 | 2 |
| CO | Cajamarca, Arenillal, Mazatlán | Tolima | 4.389 | -75.484 | 0 | 2 |
| CO | Cajamarca, La Esperanza | Tolima | 4.440 | -75.403 | 0 | 2 |
| CO | Cajamarca, San Lorenzo Alto, Los Geranios | Tolima | 4.447 | -75.396 | 0 | 2 |
| CO | Líbano, Santa Bárbara, Ambato | Tolima | 4.944 | -75.010 | 0 | 2 |
| CO | Darién, La Playa | Valle del Cauca | 3.959 | -76.461 | 0 | 2 |
| CO | Darién, La Unión, La Isabela | Valle del Cauca | 3.956 | -76.469 | 0 | 2 |
| CO | Darién, La Unión, La Violeta | Valle del Cauca | 3.925 | -76.494 | 0 | 2 |
| CO | La Unión, Córcega, Berlín | Valle del Cauca | 4.534 | -76.105 | 0 | 2 |
| CO | Palmira, Corpoica | Valle del Cauca | 3.517 | -76.300 | 0 | 37 |
| CR | | Cartago | 9.831 | -83.563 | 0 | 29 |
| CR | | Cartago | 9.784 | -83.751 | 0 | 29 |
| CU | Topes de Collantes | Sancti Spiritus | 21.856 | -79.990 | 0 | 41 |
| EC | Caluma | Bolívar | -1.631 | -79.258 | 1 | 42 |
| EC | Chillanes | Bolívar | -1.944 | -79.066 | 1 | 42 |
| EC | Rioverde | Carchi | 0.837 | -78.372 | 0 | 2 |
| EC | | Carchi | 0.840 | -78.376 | 0 | 43 |
| EC | La Maná | Cotopaxi | -0.941 | -79.232 | 1 | 42 |
| EC | Peñaherrera | Imbabura | 0.350 | -78.535 | 1 | 42 |
| EC | Garcia Moreno | Imbabura | 0.264 | -78.646 | 1 | 42 |
| EC | Palora | Morona Santiago | -1.699 | -77.962 | 1 | 42 |
| EC | Sucua | Morona Santiago | -2.456 | -78.166 | 1 | 42 |
| EC | Gualaquiza | Morona Santiago | -3.406 | -78.572 | 1 | 42 |
| EC | | Morona Santiago | -1.460 | -78.143 | 0 | 43 |
| EC | AguaYaku, Salazar | Napo | -0.897 | -77.770 | 0 | 2 |
| EC | Cotundo, Aviles | Napo | -0.712 | -77.585 | 0 | 2 |
| EC | El Chaco, Llerena | Napo | -0.406 | -77.838 | 0 | 2 |
| EC | El Chaco, Sarrias | Napo | -0.359 | -77.811 | 0 | 2 |

| Country | Locality | State/Province | Latitude | Longitude | Approx. | Reference |
|---|---|---|---|---|---|---|
| EC | Guagua Sumaco | Napo | -0.736 | -77.561 | 0 | 2 |
| EC | | Napo | -0.366 | -77.817 | 0 | 43 |
| EC | | Napo | -0.904 | -77.771 | 0 | 43 |
| EC | Puyo | Pastaza | -1.494 | -77.999 | 1 | 42 |
| EC | | Pastaza | -1.673 | -77.960 | 0 | 43 |
| EC | Pomona | Pastaza | -1.520 | -77.371 | 0 | 44 |
| EC | Los Bancos, Dr Chang | Pichincha | -0.013 | -78.890 | 0 | 2 |
| EC | Los Bancos, San Martín | Pichincha | -0.100 | -79.013 | 0 | 2 |
| EC | Tandapi | Pichincha | -0.414 | -78.799 | 1 | 42 |
| EC | | Pichincha | -0.023 | -78.894 | 0 | 43 |
| EC | Saloya | Pichincha | -0.313 | -78.710 | 0 | 44 |
| EC | Cutuglagua, Mejia | Pichincha | -0.366 | -78.550 | 0 | 45 |
| EC | El Reventador | Sucumbíos | -0.035 | -77.529 | 1 | 42 |
| EC | Gonzalo Pizarro | Sucumbíos | 0.016 | -77.378 | 1 | 42 |
| EC | | Tungurahua | -1.426 | -78.203 | 0 | 43 |
| EC | Zamora | Zamora Chinchipe | -4.062 | -78.950 | 1 | 42 |
| GF | | | 4.098 | -52.680 | 0 | 29 |
| GF | | | 4.536 | -52.123 | 0 | 29 |
| GT | | Jalapa | 14.597 | -89.983 | 0 | 29 |
| GT | Suchitepe-quez, Santa Barbara Ref Quetzal UVG | | 14.541 | -91.197 | 0 | 29 |
| HN | Comayagua | | 14.460 | -87.650 | 0 | 2 |
| HO | Pico Bonito, Estación CURLA | | 15.697 | -86.901 | 0 | 46 |
| HO | Valle de Comayagua | | 14.336 | -87.666 | 1 | 47 |
| MX | | Jalisco | 20.337 | -105.345 | 0 | 29 |
| MX | | Oaxaca | 15.926 | -96.418 | 0 | 29 |
| MX | | Veracruz | 19.520 | -96.942 | 0 | 29 |
| MX | Misantla | Veracruz | 19.925 | -96.851 | 0 | 29 |
| MX | Orizaba | Veracruz | 18.848 | -97.105 | 0 | 29 |
| MX | Presidio | Veracruz | 19.068 | -96.973 | 0 | 29 |
| PE | La Universidad Nacional Agraria de la Selva | | -9.286 | -75.998 | 0 | 48 |
| PE | Nauto | | -4.504 | -73.583 | 1 | 49 |
| PE | Loreto | | -5.066 | -73.851 | 1 | 49 |
| PE | Tingo María, Huánuco | | -9.896 | -76.279 | 1 | 49 |
| SR | Calcutta | Saramacca | 5.833 | -55.745 | 1 | 50 |
| SR | | Sipaliwini | 5.067 | -54.441 | 0 | 29 |

| Country | Locality | State/Province | Latitude | Longitude | Approx. | Reference |
|---|---|---|---|---|---|---|
| TT | Curepe, Trinidad, W.I. | | 10.630 | -61.400 | 0 | 29 |
| TT | Morne Bleu Textel Installation, Trinidad | | 10.720 | -61.290 | 0 | 29 |
| TT | Palmiste, Trinidad, W.I. | | 10.240 | -61.450 | 0 | 29 |
| VE | Pao de Zarate | Aragua | 10.114 | -67.248 | 1 | 51 |
| VE | Villa de Cura | Aragua | 10.033 | -67.493 | 1 | 52 |
| VE | Valle del Turbio, Iribarren | Lara | 10.066 | -69.333 | 0 | 2 |
| VE | El Tocuyo | Lara | 9.783 | -69.790 | 1 | 52 |
| VE | El Cuji, Barquisimeto | Lara | 10.156 | -69.306 | 1 | 53 |
| VE | El Molino, Jimenez | Lara | 9.849 | -69.613 | 1 | 54 |
| VE | la Depresion de Quibor | Lara | 9.933 | -69.609 | 1 | 55 |
| VE | San Juan de Lagunillas | Merida | 8.499 | -71.347 | 1 | 56 |
| VE | El Corozo, Caripe | Monagas | 9.694 | -63.368 | 1 | 57 |
| VE | Cordero | Táchira | 7.855 | -72.180 | 1 | 58 |
| VE | Los Rios, Seboruco | Táchira | 8.121 | -72.157 | 0 | 59 |

**S2 Table**. **Comparison of DDRP predictions for the dates of first spring egg laying ("Predicted") to the month (data set 1) or dates (data sets 2 and 3) of peak spring adult flight for *Epiphyas postvittana* in California according to three monitoring data sets ("Observed").** For monitoring data sets 1 and 2, DDRP predictions for each region are averages (and range) of several grid cells (see S2 Appendix). For data sets 2 and 3, the difference in days ("Diff") between DDRP predictions and dates of peak spring flight are presented.

| Year | Data set | Region | Observed | Predicted | Min | Max | Diff | Timing |
|---|---|---|---|---|---|---|---|---|
| 2008 | 1 | Alameda Co. | May | 4/01 | 3/27 | 4/07 | | Early |
| | | Contra Costa Co. | May | 4/15 | 3/27 | 5/19 | | In range |
| | | Monterey Co. | Apr | 4/12 | 3/17 | 5/13 | | In range |
| | | San Francisco Co. | May | 4/12 | 3/30 | 5/14 | | In range |
| 2009 | 1 | Alameda Co. | Apr | 3/21 | 3/19 | 3/26 | | Early |
| | | Contra Costa Co. | May | 4/06 | 3/18 | 5/10 | | In range |
| | | Monterey Co. | May | 4/02 | 3/02 | 5/05 | | In range |
| | | San Francisco Co. | Apr | 4/07 | 3/21 | 5/10 | | In range |
| 2012 | 2 | Region 2 | 3/01 | 3/23 | 3/19 | 4/17 | 22 | |
| | | Region 3 | 3/01 | 3/23 | 3/17 | 4/10 | 22 | |
| | | Region 4 | 3/01 | 3/22 | 3/21 | 3/24 | 21 | |
| | | Region 5 | 3/01 | 3/26 | 3/21 | 3/30 | 25 | |
| | | | | | | Average | 22.5 | Late |
| 2013 | 2 | Region 1 | 4/10 | 4/10 | 3/27 | 4/25 | 0 | |
| | | Region 2 | 4/10 | 3/29 | 3/27 | 4/08 | −12 | |
| | | Region 3 | 4/10 | 4/05 | 3/27 | 4/25 | −5 | |
| | | Region 4 | 4/10 | 3/31 | 3/30 | 4/01 | −10 | |
| | | Region 5 | 4/10 | 4/09 | 3/31 | 4/14 | −1 | |
| | | | | | | Average | −5.6 | Early |
| 2014 | 2 | Region 3 | 3/03 | 3/06 | 3/01 | 3/17 | 3 | |
| | | Region 5 | 3/03 | 3/07 | 3/02 | 3/09 | 4 | |
| | | | | | | Average | 3.5 | Late |
| 2019 | 3 | Salinas | 3/28 | 3/23 | | | −5 | Early |
| 2020 | 3 | Salinas | 3/13 | 3/15 | | | 2 | Late |

**S3 Table. DDRP predictions of the number of degree-days Celsius (DDC) that accumulated between peaks in last fall flight and first spring flight of *Epiphyas postvittana* in California.** For monitoring data set 2, predictions for each region are averages of four DDRP grid cells (see S2 Appendix). Missing results for some regions are due to indiscernible peaks in flight for one or both seasons.

| Years | Data set | Region | Fall peak | Spring peak | | DDC |
|---|---|---|---|---|---|---|
| 2011–2012 | 2 | Region 2 | 10/13 | 3/01 | | 695 |
| | | Region 3 | 10/13 | 3/01 | | 689 |
| | | Region 4 | 10/13 | 3/01 | | 705 |
| | | Region 5 | 10/13 | 3/01 | | 684 |
| | | | | | Average | 693 |
| 2012–2013 | 2 | Region 1 | 10/06 | 4/10 | | 848 |
| | | Region 2 | 11/06 | 4/10 | | 688 |
| | | Region 3 | 10/21 | 4/10 | | 758 |
| | | Region 4 | 10/06 | 4/10 | | 977 |
| | | Region 5 | 10/06 | 4/10 | | 866 |
| | | | | | Average | 827 |
| 2013–2014 | 2 | Region 3 | 10/13 | 3/03 | | 759 |
| | | Region 5 | 10/13 | 3/03 | | 728 |
| | | | | | Average | 744 |
| 2019–2020 | 3 | Salinas | 10/18 | 3/15 | | 823 |

**S1 Fig. Predictions of climatic suitability for *Epiphyas postvittana* in Australia, New Zealand, and California based on 1961–1990 climate normals according to (A, B) CLIMEX and (C) DDRP.** Climatic suitability of an area in CLIMEX is represented by the Ecoclimatic Index (EI), where EI = 0 indicates unsuitable conditions. In DDRP, the potential for long-term establishment is indicated by areas not under moderate or severe climate stress. Blue triangles in Australia and New Zealand depict the locations of training locality records used to fit the CLIMEX model (A). Black circles in California depict the locations of records used to validate the CLIMEX and DDRP models (B and C, respectively). CLIMEX maps were generated for this study and have not been previously published.

**(A)**

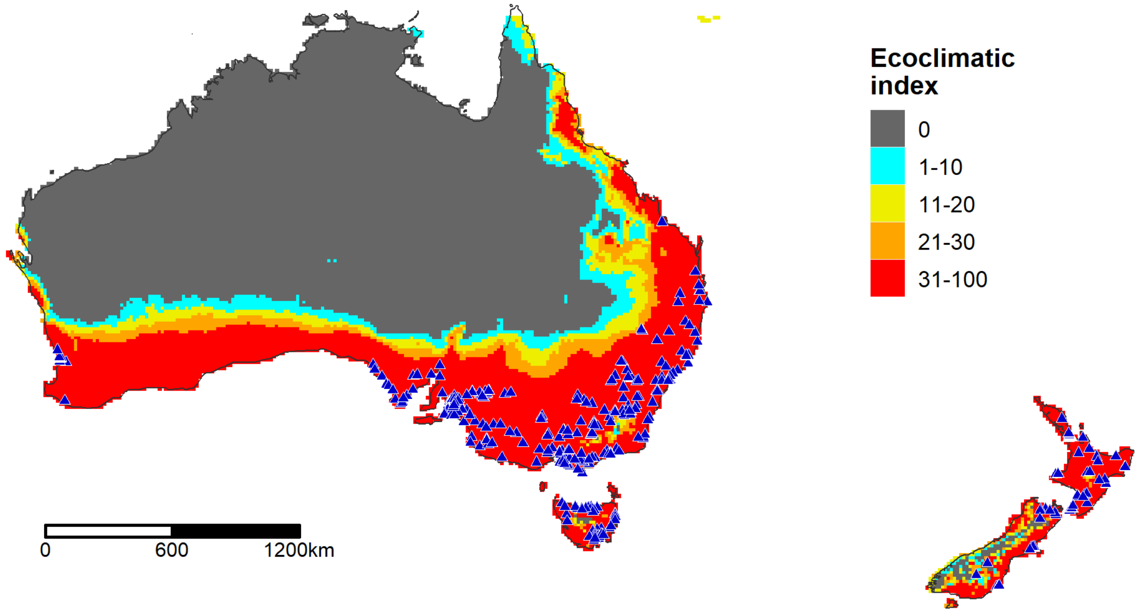

**(B)**

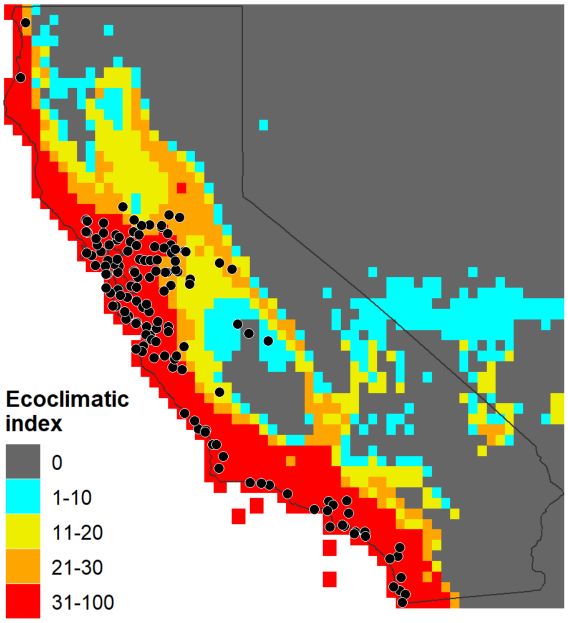

**(C)**

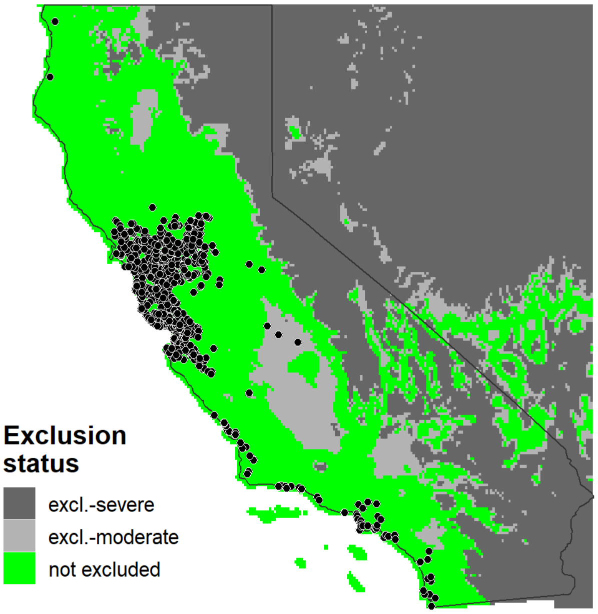

**S2 Fig. CLIMEX model for *Neoleucinodes elegantalis* in the Neotropics.** Climatic suitability of an area is represented by the Ecoclimatic Index (EI), where EI = 0 indicates unsuitable conditions. Blue triangles and black circles depict the approximate locations of occurrence records used to fit and validate the model, respectively. The map was generated for this study and has not been previously published.

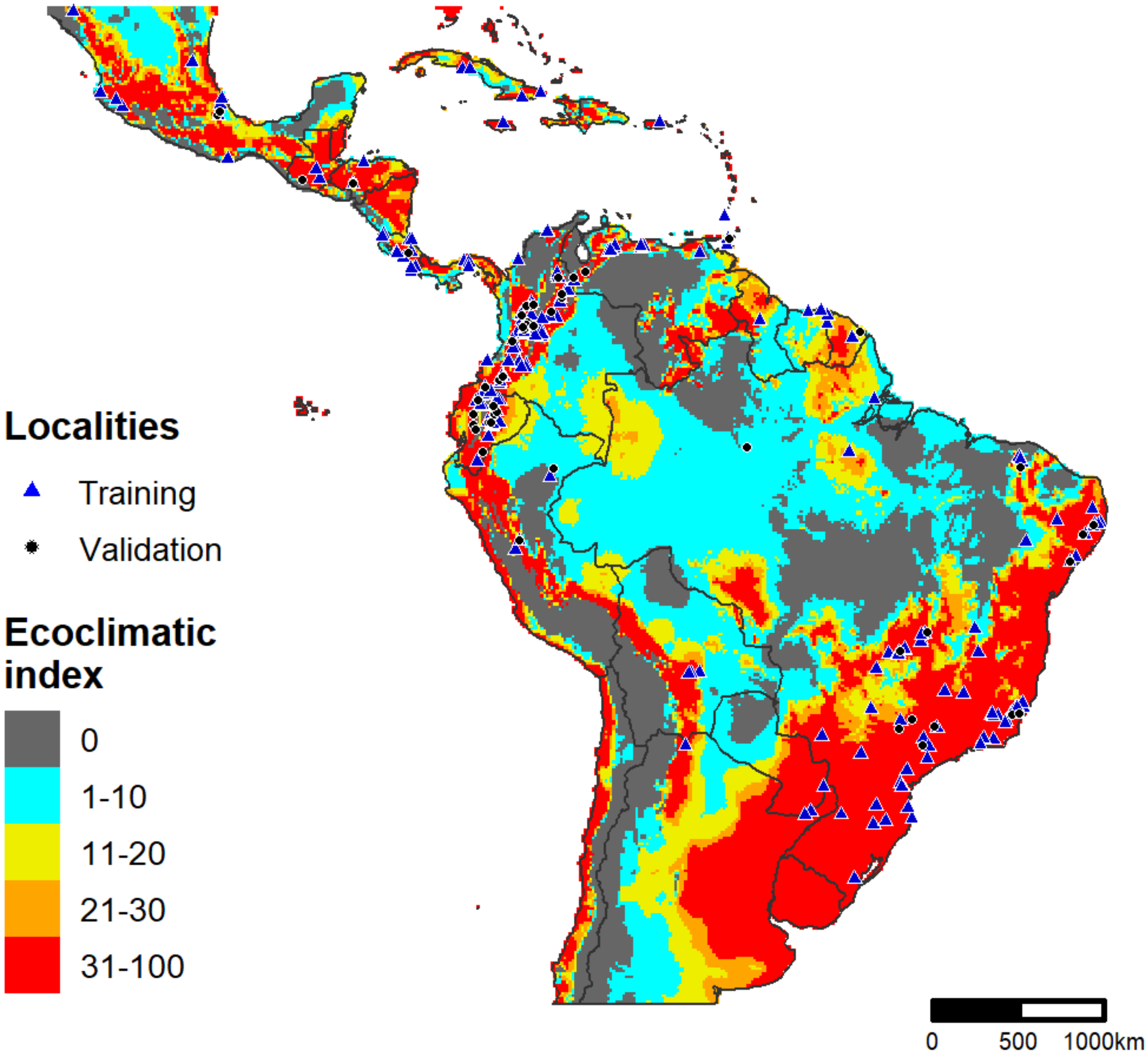

**S3 Fig. DDRP predictions of cold and heat stress for (A) *Epiphyas postvittana* and (B) *Neoleucinodes elegantalis* for 2018.** Pink and blue lines depict the moderate and severe temperature stress limits, respectively.

**(A) *Epiphyas postvittana***

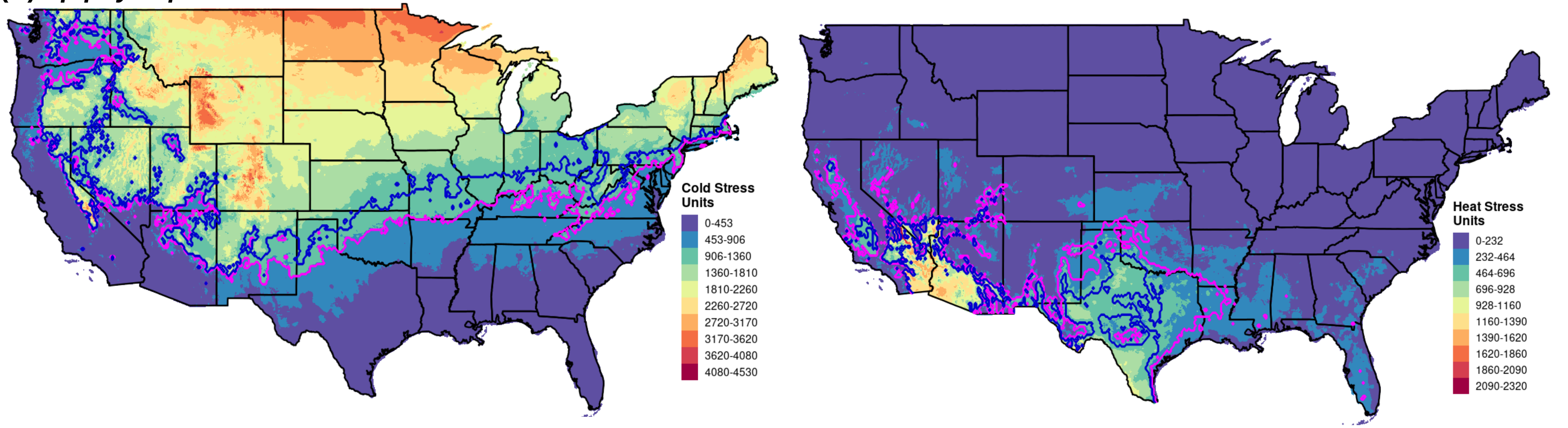

**(B) *Neoleucinodes elegantalis***

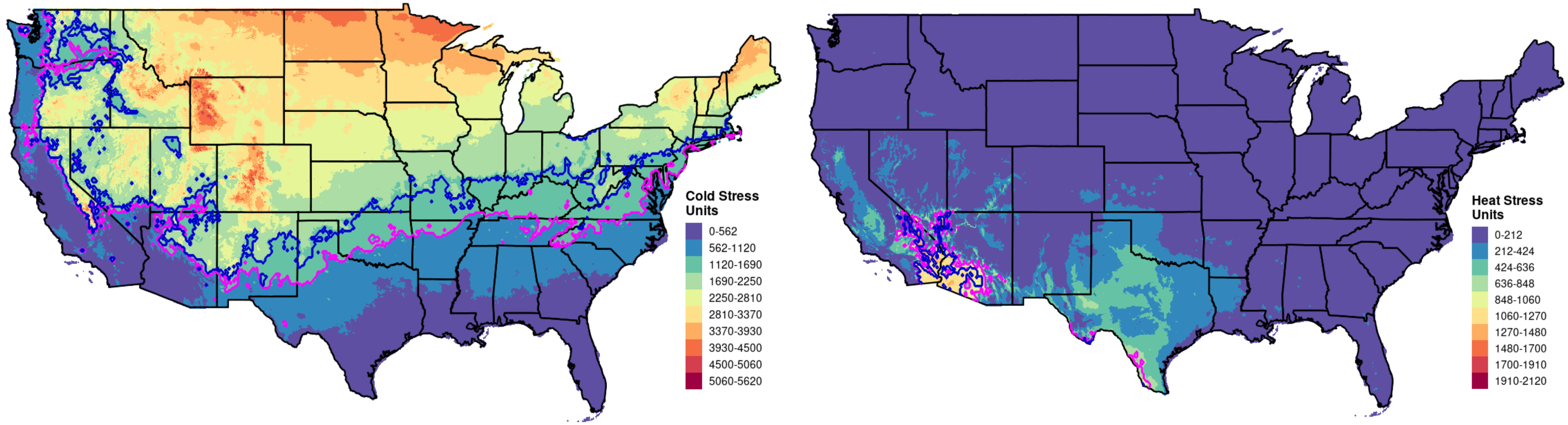

**S4 Fig. CLIMEX predictions of dry stress for *Neoleucinodes elegantalis* in CONUS.**

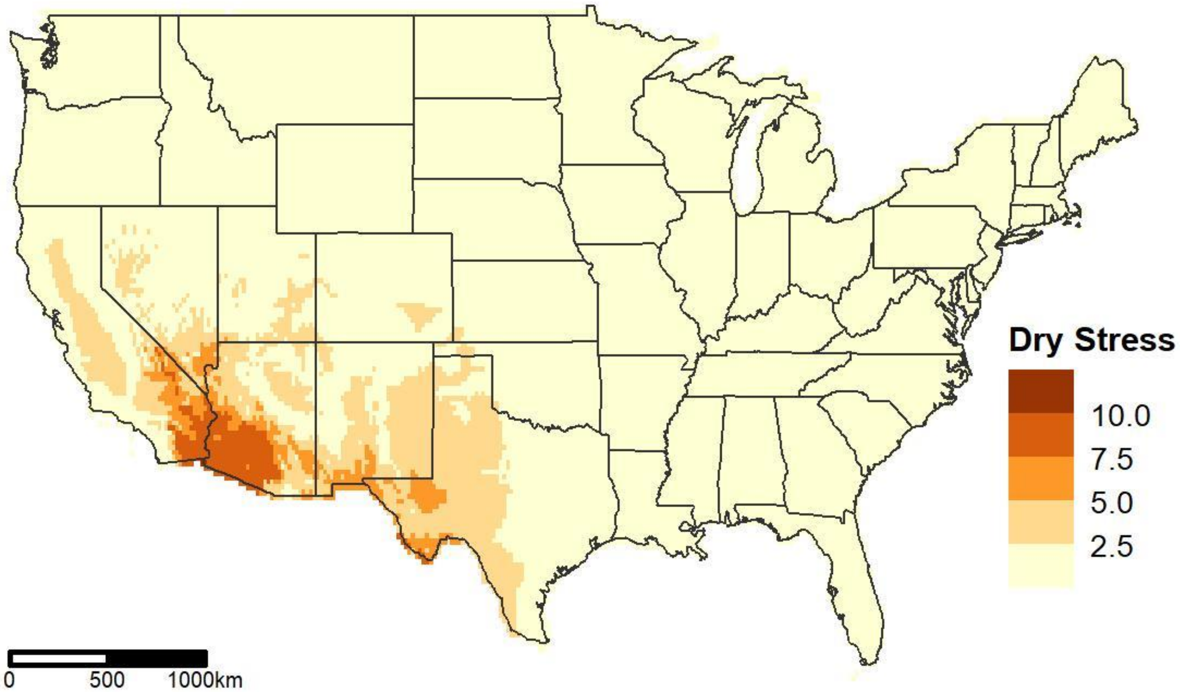
